# Respiratory vocal coordination increases as zebra finches prepare to sing

**DOI:** 10.1101/2025.05.14.654126

**Authors:** Devika Babu Kizhakoot, Shikha Kalra, PR Rubna, Sonam Chorol, Shivam Chitnis, Divya Rao, Raghav Rajan

## Abstract

Complex vocalizations like human speech and bird song require precise coordination between respiration and vocalizations with most vocalization being produced during the exhalation phase of respiration. However, when this coordination is established each time animals vocalize remains poorly understood. Here, we addressed this question by recording respiratory pressure in adult, male, zebra finches during singing. Zebra finches begin their song bouts by repeating a short vocalization called an introductory note (IN), before producing song and these INs have been hypothesized to reflect motor preparation for song. We found that each IN is associated with a large amplitude expiratory pulse. Birds that did not produce INs had a few silent, large amplitude, IN-like, expiratory pulses just before starting song. Expiratory pressure and inspiratory pressure increased with successive repeats of the IN (silent or vocalized). Further, for birds with INs, coordination between respiration and vocalization increased with vocalizations beginning earlier in the exhalation and ending later in the exhalation, such that more of the expiratory pulse was filled with the vocalizaton. Overall, these results show that respiratory vocal coordination improves with each repetition of the IN and suggest that INs reflect preparation that involves coordination of respiratory and vocal neural circuitry.

---

Most animals, including humans, generate sound by modulating the configuration and elasticity of their vocal organ and providing it with vibrational energy via respiration. Specifically, we use the expiratory air-stream from our lungs to vibrate our vocal folds [1], [2], [3]. Therefore, our vocal and respiratory apparatuses must work in tandem to facilitate speech production. Any disruption preventing this synchronization might lead to speech impairment as seen in conditions like Parkinson’s disease (PD)[4]. Since vocalization integrates various motor systems and requires their enhanced control, singing therapy is being explored to improve respiratory, vocal, and swallowing functions in PD cases [5], [6], [7]. Thus, understanding various aspects of vocal-respiratory coordination has wide-ranging implications.

How the brain prepares to coordinate respiratory and vocal circuitry, to produce vocalizations, remains poorly understood. The vocal apparatus has been studied in isolation in cue-triggered tasks and motor preparation before vocal onset has been observed[8]. However, how coordination is established during actual vocalizations and during self-initiated vocalizations remains unclear.

Like humans, songbirds use similar biomechanics to generate vocalizations through their vocal organ, the syrinx [9], [10]. Adult male zebra finches (a well-studied Australian songbird) produce complex vocalizations (songs), as part of their courtship display. They also produce these songs when they are alone. Thus, songbirds become a good model to study motor preparation before self-initiated vocalizations. These birds typically start their songs with short repeated elements called introductory notes (INs)[11]. The acoustic properties of INs and the associated neural activity show signatures of motor preparation[12], but how respiration changes during these INs is unclear. Here, we investigated respiratory patterns during INs and their alignment with the vocal output to understand how birds coordinate vocal and respiratory motor systems at the onset of singing.

## Analysis of Respiratory Pressure Patterns During Introductory Notes

We cannulated the air sac to record respiratory pressure during singing in 10 adult male zebra finches (Fig. 1B, Fig. S1A). As in previous studies, respiratory patterns during singing were distinct from those during silence [13]**(Fig. 1A)**. Each unit of vocalization or syllable occurred within a single expiration (mean = 1.02, SD = 0.13, n = 5100 expiratory pulses across 10 birds, range = 1.000 to 1.084, Table S1 and Figure S1), while each inter-syllable interval was marked by a single deep inspiration.

**Fig 1.**
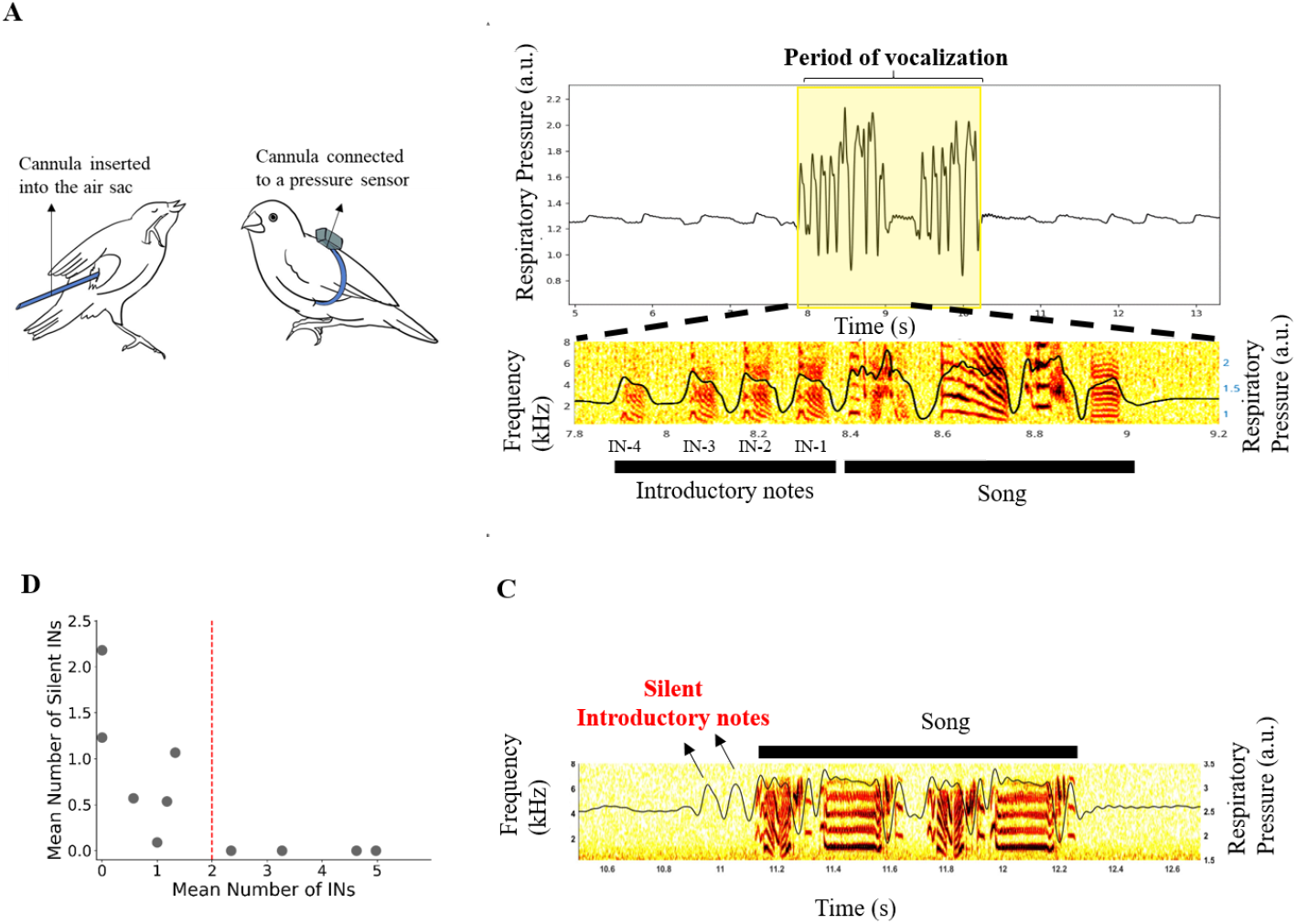
High amplitude expiratory pulses are produced without corresponding vocalization at onset of song in low IN birds. (A) A schematic of air sac pressure recording. A cannula is inserted into the air sac of the bird and connected to a pressure sensor to record respiratory pressure patterns. (B) Represented is the simultaneously recorded air-sac pressure and sound. Air sac pressure pattern of a bird during quiet respiration (portions flanking the yellow box on either side) and vocal respiration (yellow box) (bottom). The corresponding spectrogram during a segment of song bout is overlaid with respiratory pressure (black) (bottom). The song has 4 INs preceding it marked as IN-4 to IN-1. (C) A segment of song sung by a bird which does not produce any INs. Air sac pressure pattern associated with the song (black trace) is overlaid upon the song spectrogram. Arrows point to high amplitude expiratory pulses which precede the song, called Silent INs. (D) Mean number of INs produced by the bird Vs Mean number of Silent INs the bird produces. All birds with mean IN < 2 (red line) produce Silent INs.

Zebra finches sing a stereotyped song, in which the motif sequences remain consistent across multiple renditions. However, these motifs are preceded by a variable number of Introductory Notes (INs). INs have been proposed to reflect preparatory motor activity in the brain prior to song initiation [12].Contradicting this assumption, some zebra finches are known to produce songs without any preceding INs. To investigate this, we examined the respiratory pressure patterns associated with songs in two birds that consistently sang without INs (Fig. 1C). Notably, these birds initiated their songs with high-amplitude expiratory pulses, despite the absence of audible vocalizations. We refer to these non-vocal preparatory events as **Silent INs** (Fig. 1C). Similar to vocalized INs, the number of Silent INs increased when these birds sang directed songs in the presence of a female.

To assess whether Silent INs are unique to these individuals, we analyzed recordings from additional birds. We found that those producing few vocal INs before song often exhibited Silent INs, whereas birds that produced more than two INs on average did not (Fig. 1D).

Since Silent INs share key features with vocalized INs, we will treat them equivalently in our analyses, except when explicitly examining respiration in relation to vocal output.

## Amplitude of Expiration and Inspiration increases as Introductory Notes progress

One characteristic feature of transitioning from quiet respiration to vocalization is an increase in expiratory pressure as it is essential to attain subglottal pressure necessary for speech production. Therefore expiratory pressure is correlated with the two behavioural states of the bird: quiet vs singing. But upon examining expiratory amplitudes (Figure 2A) during quiet respiration, during INs and during the first syllable of song motif, INs marked a distinct transition state-from the low expiratory amplitudes corresponding to quiet respiration and the high expiratory pressure during song (Figure 2B, left).

**Fig 2.**
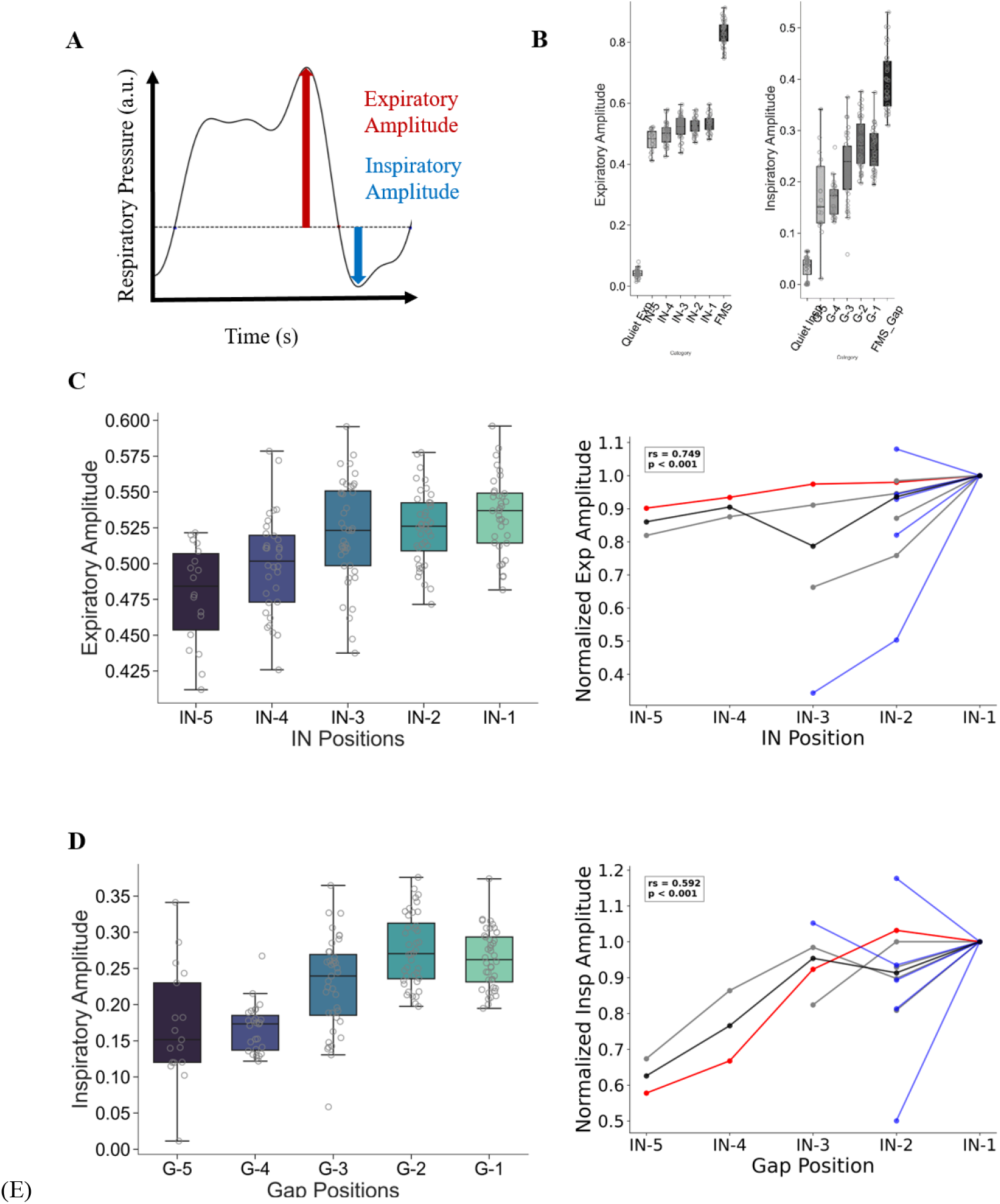
Expiratory and Inspiratory amplitudes increase as INs progress. (A) A schematic diagram of parameters of air sac pressure (black trace) used in the analysis. Expiratory peak (red arrow) is the highest pressure value within an expiratory pulse and its height from the baseline (black dotted line) is called expiratory amplitude. Inspiratory peak (blue arrow)is the lowest pressure value within an inspiration and its height from the baseline is called Inspiratory amplitude. (B) Expiratory Amplitudes corresponding to Quiet expiration, different IN positions and first syllable of the motif (FMS) for one example bird (left). Inspiratory Amplitudes corresponding to Quiet inspiration, different Gap positions and the inspiration between first and second motif syllables (FMS_Gap), for one example bird (right). (C) Expiratory Amplitudes across different IN positions for one example bird, left. Median expiratory amplitudes are positively correlated with IN positions (n =10 birds), right. Median at each position is normalized to the value at the last IN.Grey lines represent individual birds, blue lines represent birds that produce silent INs, the red line represents the example bird and black line represents median across birds. (D) Inspiratory Amplitudes across different IN positions for one example bird, left. Median inspiratory amplitudes are positively correlated with IN positions (n =10 birds), right. Median at each position is normalized to the value at the IN-1. Grey lines represent individual birds, blue lines represent birds that produce silent INs, the red line represents the example bird and black line represents the median across birds.

Higher amplitude songs produced by birds are correlated with higher sub-syringeal air sac pressure [14]. And INs progressively get louder from the first one produced to the last one produced before motif onset [12]. Therefore, we wanted to extend these two observations to understand how expiratory amplitudes (Figure 2A) change as INs progress. We calculated the expiratory amplitude (Figure 2C) value corresponding to each IN and found that it increases as the sequence progressed towards the last IN, which we call IN_-1_ (ρ= 0.749, p < 0.001 for n = 10 birds, Spearman’s rank-order correlation,Figure 2C).

While the bird sings, it takes short inhalations in the gaps between syllables. These inhalations are deeper or of higher amplitude compared to those during quiet breathing (Figure 2B, right). Since we had found expiratory amplitudes increase as INs progress we next asked if inspiration in the gaps between INs also follow a similar pattern. Inspiratory amplitudes (Figure 2A) were calculated by taking the lowest pressure value corresponding to each Gap. We found that birds take deeper inspirations as the sequence progressed towards the gap between last IN and the beginning of motif, which we call Gap_-1_ (ρ= 0.592 p < 0.001 for n = 5 birds, Spearman’s rank-order correlation, Figure 2D)

## Vocalization and respiration get better coordinated as Introductory Notes progress

Previous work shows that as zebra finches mature from juvenile to adult, their vocal-respiratory coordination improves: a greater proportion of expirations gets filled by a corresponding syllable [15].Therefore in adult zebra finches there is a strong vocal-respiratory coordination during song. But within adult birds we wanted to investigate if such coordination is achieved during song by using INs to bring about progressive improvement. We used coordination ratio (syllable duration/ expiratory pulse duration) (Figure 3A) as a measure of such coordination. We found that coordination ratio increases as the sequence progressed towards the last IN and gets closer to the coordination achieved during the first motif syllable (ρ= 0.856, p < 0.001 for n = 5 birds, Spearman’s rank-order correlation, Figure 3B).

**Fig 3.**
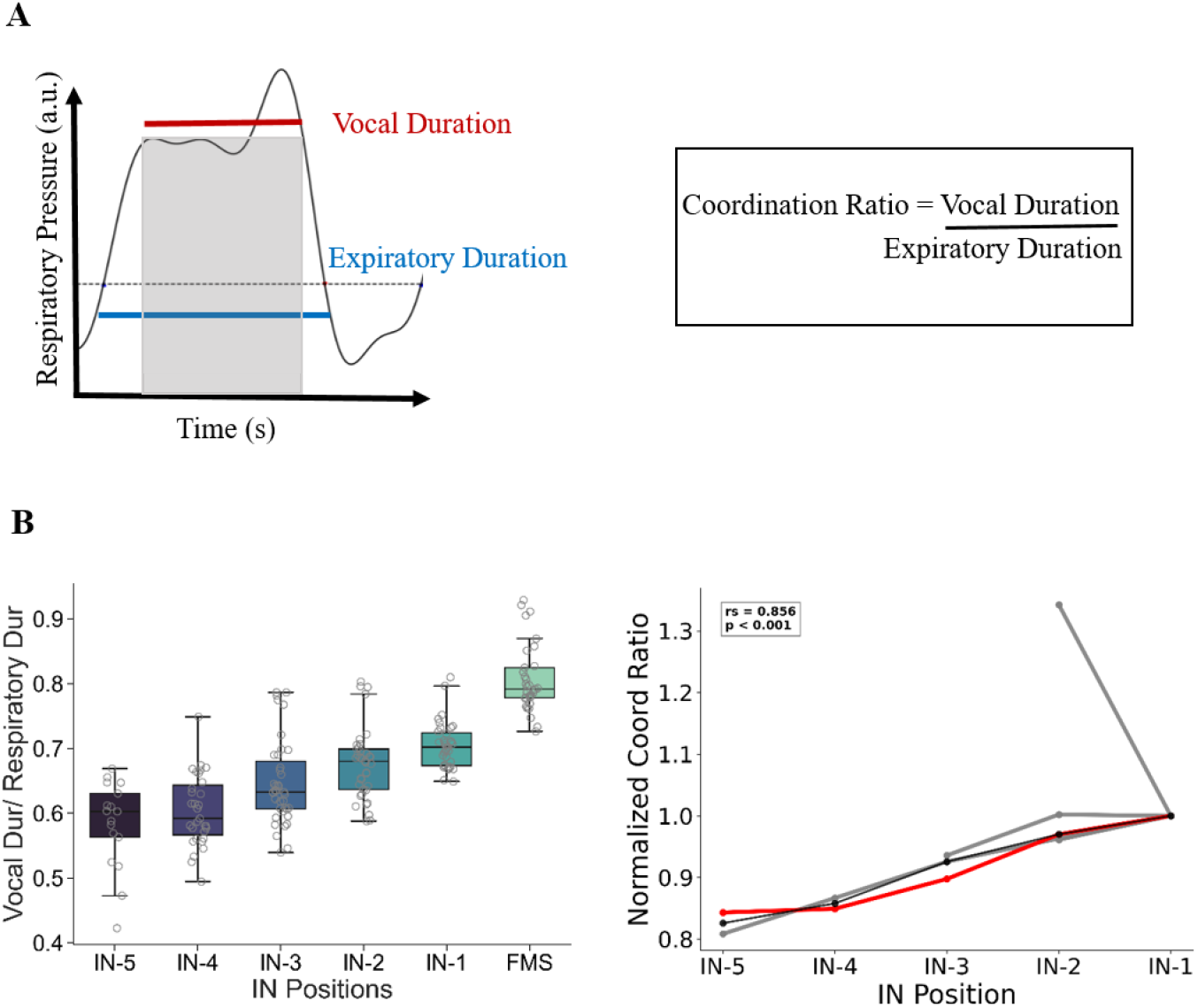
Vocal-Respiratory Coordination improves as INs progress. (A) A schematic diagram of parameters of air sac pressure (black trace) (left) used in the analysis and the formula used (right). Grey Bar represents a syllable sung within the expiratory pulse. Vocal duration is the duration of that syllable and respiratory duration is the duration of the corresponding expiratory pulse. Respiratory duration extends from the expiratory onset to expiratory offset. (B) Coordination ratio across different IN positions for one example bird, left. Median coordination ratio values are positively correlated with IN position (n =5 birds), right. Median at each position is normalized to the value at the last IN.Grey lines represent individual birds, the red line represent the example bird and black line represents median across birds.

**Fig 4.**
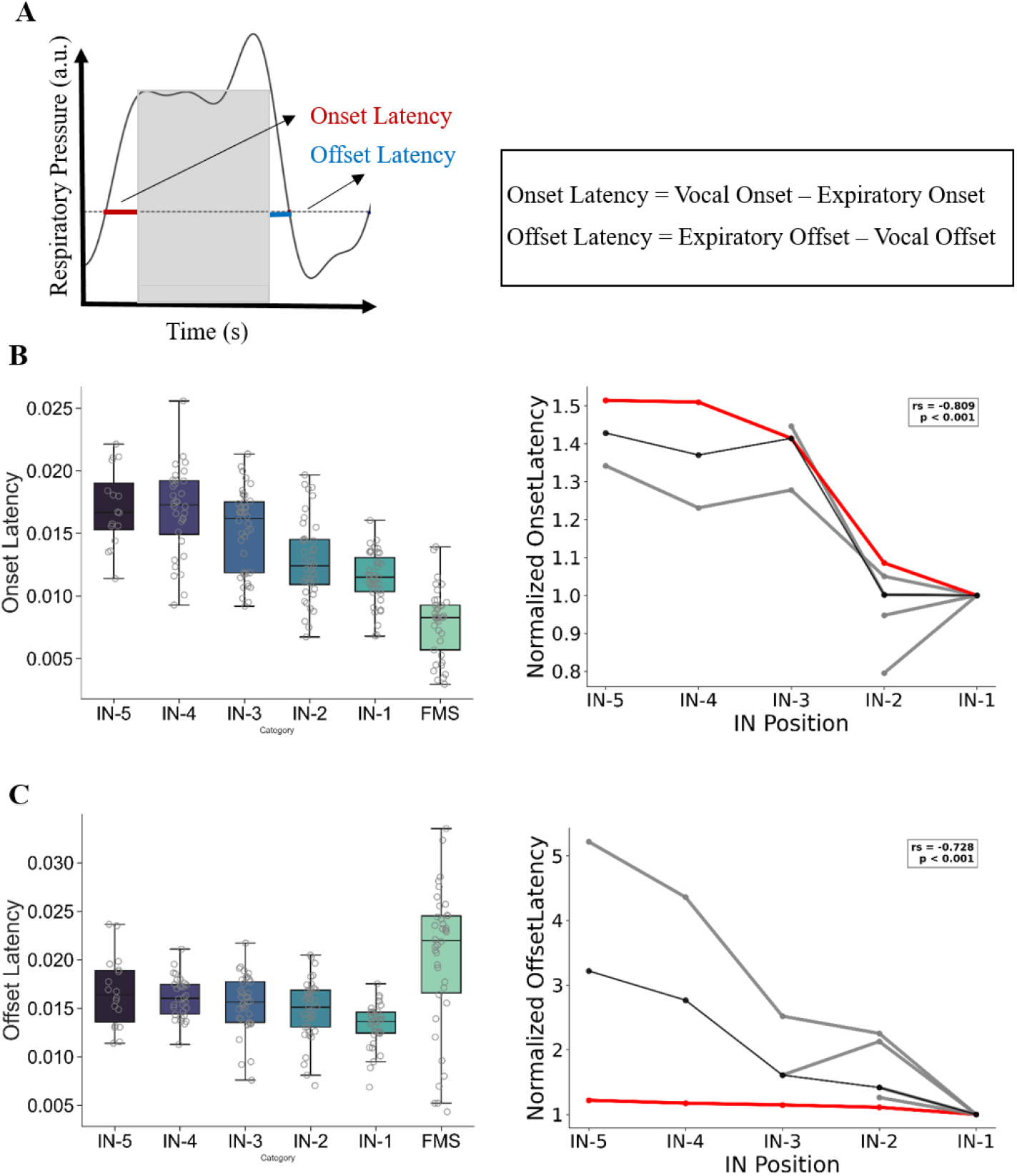
Expiration starts and ends closer to the vocal output as INs progress. (A) A schematic diagram of parameters of air sac pressure (black trace) (left) used in the analysis and the formula used(right). Grey Bar represents a syllable sung within the expiratory pulse. Onset latency is the duration between expiratory and vocal onsets. Offset latency is the duration between vocal and respiratory offsets. (B) Onset latency across different IN positions for one example bird, left. Median Onset latency values are negatively correlated with IN position (n =10 birds), right. Median at each position is normalized to the value at the last IN. Grey lines represent individual birds, the red line represents the example bird and black line represents the median across birds. (C) Offset latency across different IN positions for one example bird, left. Median offset latency values are negatively correlated with IN position (n =5 birds), right. Median at each position is normalized to the value at the last IN. Grey lines represent individual birds, the red line represents the example bird and black line represents the median across birds.

We further analysed other parameters to understand such a trend in coordination ratio. We found that this could be explained by a decrease in the duration between the start of expiratory pulse and start of vocalization: onset latency and a decrease in the duration between the end of vocalization and expiration: offset latency (Figure 3A). Both decreased as INs progressed towards song (for onset latency,ρ= −0.809, and for offset latency,ρ= −0.728, p < 0.001 for n = 5 birds, Spearman’s rank-order correlation, Figure 3B and 3C)

## Discussion

Vocalization is a complex motor behavior that requires the coordination of multiple motor systems to generate sound. In particular, with respect to the respiratory system—which is typically regulated by brainstem centers—vocalization likely reflects a shift in control to cortical regions involved in speech and language [16]. Similarly in birds during vocalization, the song motor pathway must be able to modulate respiration in order to generate precisely timed, high amplitude expiratory pulses that facilitate song [17]. Since these planned motor actions are preceded by neural motor preparation, and INs are thought to reflect vocal outputs of this process, they serve as useful candidates for studying how the brain prepares the respiratory and vocal systems to facilitate song.

Here, we show that IN sequences show a progressive improvement in vocal-respiratory coordination which can be explained by a similar progressive decrease in onset and offset latencies. This suggests that the muscle activation that starts and stops expiration and vocalization, under respiratory and vocal systems get better synchronized as INs progress. Since during song there is strong coordination, the bird might be using INs as a transition stage to improve vocal-respiratory coordination at the onset of singing. A similar trend is also observed in the expiratory and inspiratory amplitudes as INs progress.

In birds that initiate song with few or no audible INs, we find that high-amplitude expiratory pulses—termed *silent INs*—often precede song onset. Although the precise biomechanics underlying this phenomenon remain unclear, these silent INs may reflect the process by which respiratory centers and the song motor pathway achieve coordinated neural activity. Supporting this idea, we observe an increase in the number of silent INs when the bird sings to a female, suggesting that higher levels of coordination—or a longer time to reach an optimal preparatory state—may be required in this context. Since INs are learnt [18], it would also be interesting to understand how juvenile birds learn to not sing vocal INs and begin song with silent INs.

Overall we associate a novel function to INs as a necessary transition state to start singing. It is similar to various steps in parameter space (respiratory and vocal-respiratory) that the birds must climb before attaining levels necessary for song. Although perturbation studies are required to causally prove this hypothesis.

## Methods

All experimental procedures were done after obtaining the approval from the Institute Animal Ethics Committee (IAEC), IISER Pune and were in accordance with the guidelines of CCSEA (Committee for the Control and Supervision of Experiments on Animals), New Delhi.

Zebra finches purchased from outside vendors (n=6) or bred in IISER Pune (n=4) were used for all experiments.

### A) Air sac Cannulation Surgery

All surgeries were done under anaesthesia. 1 – 1.5 hrs before surgery birds were orally given meloxicam (0.25mg/kg), an analgesic. Surgeries were done under anesthetic isoflurane maintained at a flow rate of 1L/min and 2-4% concentration. During surgery, a heat lamp was continuously used as birds are not able to maintain body temperature under anaesthesia. Birds were laterally placed on a foam sheet and kept in place using surgical tape. A subcutaneous injection of a lignocaine (local anaesthetic) was given just below the last rib and an incision was made there. Following this, the site was punctured using forceps, and a tube was inserted. The cannula tube was ~7 cm long (inner diameter 1mm, outer diameter 2mm) and had an outer tube, 3mm long (inner diameter 2mm, outer diameter 3mm) fitted over it. Some saline was injected into the end of the cannula tube. The pulsating column of saline and the presence of vapours were used as a confirmation for detecting fluctuations in respiratory pressure. The tube is sutured to the last rib and surrounding skin. Tissue adhesive (Vetbond) is applied over that area and the bird is placed in a home cage for recovery.

### B) Air sac Pressure and Song Recording

#### Circuit Design

A pressure sensor (FMG-302PGSR) from Fujikura and an amplifier (INA 122U) made by Texas Instrumentation were used for recording respiratory pressures. The voltage and ground were connected to the Arduino board. The output was connected to the data acquisition board. The amplifier and sensor were epoxied onto a rubber band which was put on the bird as a backpack. The output from this was connected to a commutator to allow for movement of the bird.

#### Sound and Respiratory Pressure recording

One day after air sac cannulation surgery the backpack was placed on the bird.

The songs were recorded both in the absence (undirected song) of females. The songs were recorded inside a custom-made sound attenuation chamber with an omnidirectional (AKG 517; Countryman Associates, Menlo Park, CA) microphone placed at the side of the cage. Both respiratory and sound recordings were acquired simultaneously using a data acquisition board (National Instruments, NIDAQ PCI) at a sampling rate of 32000 Hz. Data was continuously recorded for a few days. This data was directly stored in the computer and later used for analysis. Signals from the pressure sensor were filtered with a 25 Hz low pass filter.

### C) Data Analysis

Custom-made scripts in Matlab (Mathworks) and Python were used for analysis.

#### Sound Data Analysis

##### Labelling and segmentation

The sound data acquired was segmented using an amplitude threshold with a minimum syllable duration of 10ms and minimum inter-syllable interval of 5ms. Song files were labeled in a semiautomated manner using a template matching algorithm which was manually reviewed later. Song bouts were defined as a sequence of vocalizations separated by 2 seconds of silence. A motif was defined as a frequently occurring sequence of syllables. repeating vocalizations other than call occurring before motif syllables were defined as introductory Notes (IN). Calls were defined as vocalizations that occurred in isolation.

##### Counting IN number

Consecutive INs occurring before the first motif syllable that are separated by less than 500ms are considered for calculating the mean IN number of a bird. A minimum 20 song bouts are used for this calculation.

#### Respiratory Data Analysis

##### Defining Exhalation Offsets and Onsets

The baseline respiration was determined by calculating the median respiratory amplitude of one - three, 1-second chunks of silent respiration separated by one second from any vocalization. The first respiratory amplitude datapoint crossing this threshold was marked as an expiratory onset while the first point at which the trace went below this threshold was marked as an expiratory offset

##### Coordination between Vocalization and Respiration

The closest expiratory onset that occurs before the onset of vocalization is the expiratory onset associated with that syllable. The closest expiratory offset that occurs after the offset of vocalization is the expiratory offset.

The coordination between vocalization and respiration was calculated as the difference between the vocal onset/offset and expiratory onset/offset.

##### Counting Silent exhalations

Non-vocalization-associated respiratory amplitude within an expiratory onset and offset was considered a silent exhalation if its peak respiratory amplitude was greater than the mean of baseline plus 6 standard deviations. Consecutive silent exhalations not separated for more than 500 ms were used for calculating the mean of silent exhalations.

##### Determining the Peak expiration and inspiration

Within a respiratory onset and respiratory offset the maximum respiratory amplitude was considered to be the peak expiration. Expiratory amplitude is taken as the difference between peak expiration and baseline respiration. Similarly, for peak inspiration the minimum respiratory amplitude after the respiratory offset was taken. Inspiratory amplitude is taken as the difference between baseline respiration and peak inspiration.

## ACKNOWLEDGMENTS

We would like to thank Prakash Raut for help with bird colony maintenance. We would like to thank Lena Veit for help with respiratory pressure recordings, Mimi Kao, Michael Long and members of the Rajan Lab for useful discussions.

