## Supplementary Figures for "Respiratory vocal coordination increases as zebra finches prepare to sing"

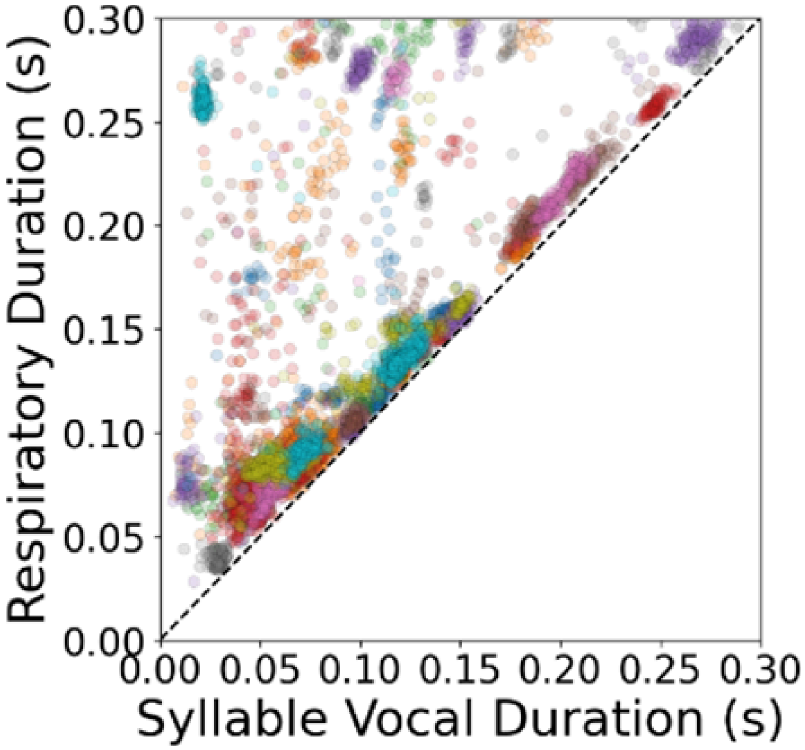


**Fig S1 | Syllables are well coordinated with corresponding respiration**

Vocal Duration vs Respiratory Duration for n=5100 syllables across n= 10 birds. Each colour represents a different bird

| Bird ID | No. of Expiratory Pulses (EP) | Mean No. of Syllables per EP | Standard Deviation |
|---|---|---|---|
| Blue121pink113 | 425 | 1.020772 | 0.048450 |
| Blue135pink127 | 674 | 1.020772 | 0.152668 |
| Green139 | 216 | 1.041667 | 0.199826 |
| Green162white171 | 719 | 1.025035 | 0.164893 |
| Green192white155 | 559 | 1.007156 | 0.084288 |
| Green194white196 | 336 | 1.029762 | 0.186624 |
| Pink030blue76 | 535 | 1.000000 | 0.000000 |
| Red084pink176 | 252 | 1.083665 | 0.276885 |
| Yellow184pink176 | 618 | 1.000000 | 0.000000 |
| yellow198pink155 | 767 | 1.002608 | 0.050998 |

**Table S1 | Only one syllable per expiratory pulse is sung**

Across n=10 birds, no of expiratory pulses are analyzed to find the number of syllables within them.
